# The Language of Cancer: Decoding Cancer Signatures with Language Models and cfRNA Sequencing

**DOI:** 10.1101/2024.06.29.601341

**Authors:** Siwei Deng, Lei Sha, Yongcheng Jin, Tianyao Zhou, Chengen Wang, Qianpu Liu, Hongjie Guo, Chengjie Xiong, Yangtao Xue, Xiaoguang Li, Yuanming Li, Yaping Gao, Mengyu Hong, Junjie Xu, Shanwen Chen, Pengyuan Wang

## Abstract

We present GeneLLM, a novel large language model that offers a transformative approach to non-invasive cancer detection and biomarker discovery by directly interpreting plasma cell-free RNA (cfRNA) sequences. Unlike traditional annotation-dependent methods, GeneLLM operates without prior knowledge, achieving significantly improved multi-cancer detection accuracy. Critically, GeneLLM identifies novel cfRNAs (‘pseudo-biomarkers’) originating from previously unannotated genomic regions–overlooked by existing methods–offering new therapeutic targets and insights into intercellular communication. This innovative, cost-effective approach bypasses traditional bioinformatics tools, generating novel ‘pseudo-biomarkers’ that outperform existing methods even with low-depth sequencing data. Consequently, GeneLLM opens new avenues for biomarker discovery and expands our understanding of the extracellular transcriptome’s role in cancer development.

## 1 Main

Early cancer detection is crucial for improving treatment efficacy and patient outcomes [9, 29]. Non-invasive, cost-effective techniques, such as those leveraging cell-free RNA (cfRNA), are increasingly important in this endeavour. CfRNA offers advantages over cell-free DNA (cfDNA) for solid tumour detection due to its ability to reflect dynamic gene expression profiles in response to disease [2, 8, 21, 26, 27, 30, 32]. This approach addresses limitations of existing methods like colonoscopy and mammography, which can be invasive, expensive, and resource-intensive [3, 5].

Nonetheless, cfRNA-based detections face challenges related to sensitivity and specificity, arising from fluctuating cfRNA levels and the difficulty in differentiating cancerous and normal profiles. To tackle this, studies have integrated machine learning methods, typically performing Differentially Expression (DE) analysis to discover Differentially Expressed Genes (DEGs) and identify potential biomarkers using methods such as random forest [24]. However, these methods are still reliant on annotated genes.

Intriguingly, research has uncovered sequences from cfRNA that do not usually reside in gene regions, such as repetitive RNA regions, which show potential as biomarkers [26]. Other transcripts, like transcribed ultra-conserved regions, also exhibit promise in cancer detection [4]. These sequences, often referred to as the genome’s ‘dark matter’, are gaining recognition in cancer detection [20]. Despite their potential, these sequences are not annotated and thus cannot be included in traditional bioinformatics pipelines dependent on genome annotations. Consequently, the valuable insights from the sequences transcribed from the genome’s ‘dark matter’ are frequently overlooked.

To harness this untapped potential, we introduce Gene Large Language Model (GeneLLM), an innovative pretrained, cfRNA-based large language model. GeneLLM directly analyses raw cfRNA sequencing data, enabling the discovery of short, cancer-indicative cfRNA sequences, termed ‘pseudo-biomarkers’, without relying on existing genome annotations. In this study, demonstrates GeneLLM’s effectively classified multiple cancer types using cfRNA samples from multiple centres, achieving consistent performance with significantly reduced sequencing depth (one-sixth of standard depth), thereby lowering sequencing and computational costs. Importantly, GeneLLM identified cfRNA molecules transcribed from unannotated regions of the genome, which are undetectable using conventional methods.

Our findings have significant implications. Firstly, GeneLLM revolutionises marker discovery by directly identifying short cfRNA sequences (‘pseudo-biomarkers’) without the need of prior knowledge. It can even identify sequences from unannotated regions (the ‘dark genome’), often overlooked by traditional methods. Secondly, GeneLLM improves cancer detection accuracy and reduces costs by requiring only plasma samples to detect multiple cancers simultaneously. Its versatility across different cancer types underscores its diagnostic potential and suggests it could revolutionise cancer screening by unlocking the potential of the ‘dark genome’, making it more accessible and affordable. Finally, our framework can be applied to other sequencing-based datasets, such as intracellular bulk RNA-seq and single-cell RNA-seq, to discover new ‘pseudo-biomarkers’ and potential therapeutic targets.

## 2 Study design and sample characteristics

CfRNA sequencing and subsequent analysis using the pretrained GeneLLM model were performed on samples from four cancer types and matched non-cancer controls, collected across three centres: Sir Run-Run Shaw Hospital, Beijing Hospital, and Peking University First Hospital (Figure 1A–E). Figure 1F and Table S1 summarise the 496 cases included in the study, with the majority of cancer samples (67.8%) originating from early-stage disease (stages I and II). Ribosomal RNA (rRNA) depletion prior to sequencing resulted in low rRNA content within the samples. The cfRNA transcripts predominantly comprised protein-coding RNA and non-coding RNA (ncRNA), including long non-coding RNA (lncRNA) but excluding rRNA (Figure 1G).

**Figure 1.**
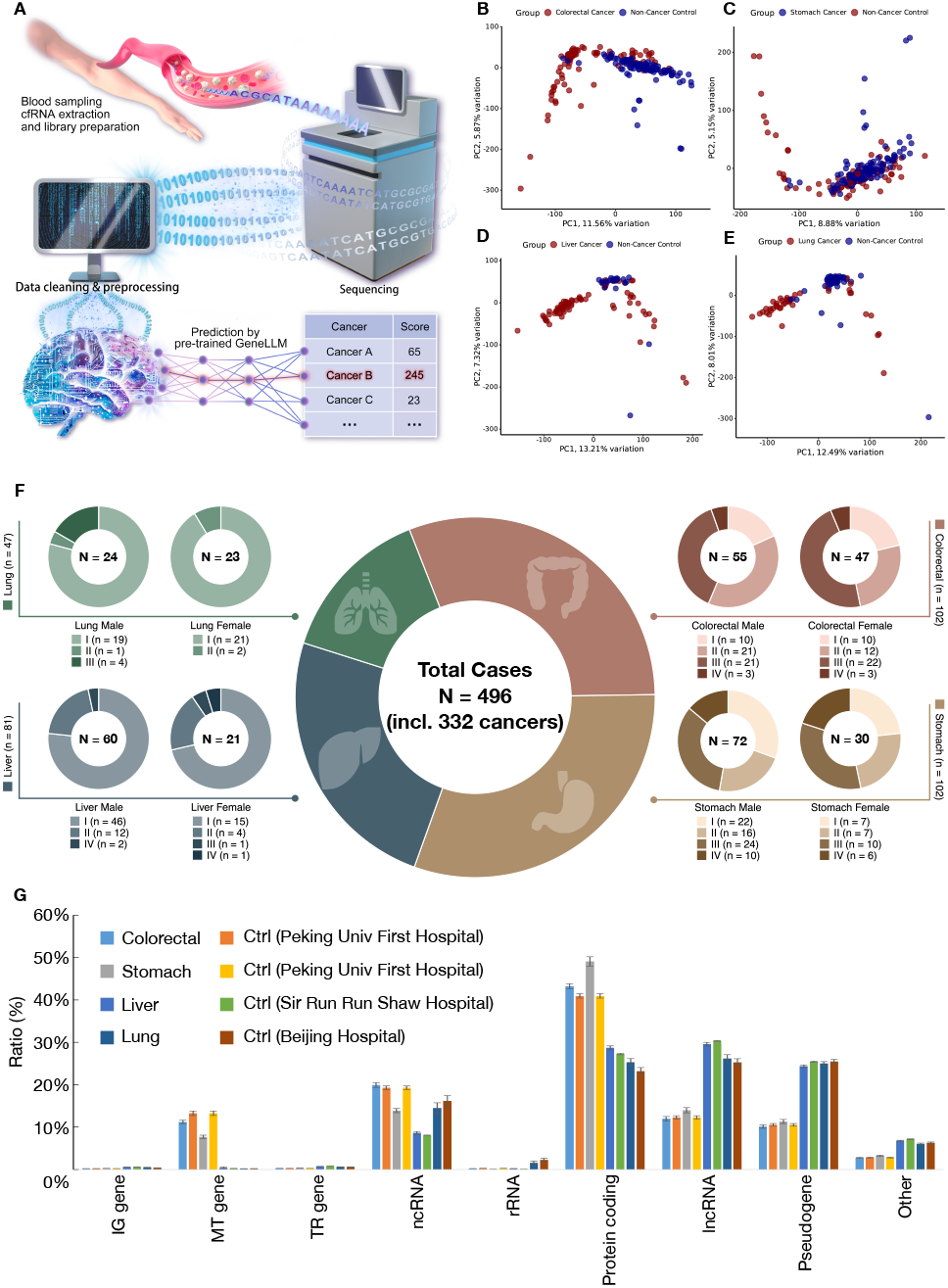
Multi-centre, multi-cancer study design. (A) Study workflow: Blood is drawn from the median cubital vein, followed by plasma centrifugation for cfRNA extraction. Libraries were prepared and sequenced using the Illumina NovaSeq 6000 platform. Cleaned sequencing reads were analysed using the pretrained large language model, GeneLLM, to predict the likelihood of each cancer type. (B–E) PCA plots showing cancer samples (dark red dots) and centre-matched non-cancer controls (dark blue dots). Non-cancer controls were matched by centre to mitigate batch effects. (F) Distribution of cancer patients by gender and stage. (G) Distribution of RNA biotypes across cancer samples and matched non-cancer controls. Abbreviations: IG gene = immunoglobulin gene; MT gene = mitochondrial gene; TR gene = T cell receptor gene; ncRNA = non-coding RNA; rRNA = ribosomal RNA; Protein coding = RNA that encodes protein-coding genes; lncRNA = long non-coding RNA; Pseudogene = RNA that encodes pseudogenes; PCA = principal component analysis

## 3 Model design and performance evaluation

### 3.1 GeneLLM’s architecture

GeneLLM leverages the sequential data processing capabilities of large language models to improve the accuracy of early cancer detection. We constructed and pretrained a cfRNA-based large language model using unlabelled cfRNA reads to establish a deep understanding of the relationships within these sequences. Given the large volume of cfRNA reads per patient (approximately 40 million), we fine-tuned GeneLLM’s cancer detection module using labelled patient data after pretraining. This module, acting as a complex language model head, extracts valuable information from the vast amount of cfRNA reads to generate cancer detection predictions.

GeneLLM’s main architecture processes each patient’s extensive cfRNA reads, compares them, and identifies minor anomalies indicating potential disease. The model operates in three main stages. First, we pretrained the Transformer architecture using cfRNA reads (Figure 2A). Since disease labels are not required at this stage, we mixed and shuffled each patient’s cfRNA reads, then tokenised them into 7-mer sequences. The pretraining process follows the same methodology as that of language models [25]. Second, once GeneLLM pretrained, GeneLLM transforms each cfRNA sequence into a vector by the output hidden layer of GeneLLM’s last output. We then combined all patients’ cfRNA vectors to mine pseudo-biomarkers, which are the most representative cfRNA sequences. Third, we constructed a disease judgement head (Figure 2B) and conducted disease tuning based on the pseudo-biomarkers. Further details regarding these stages are provided in Section 6.

**Figure 2.**
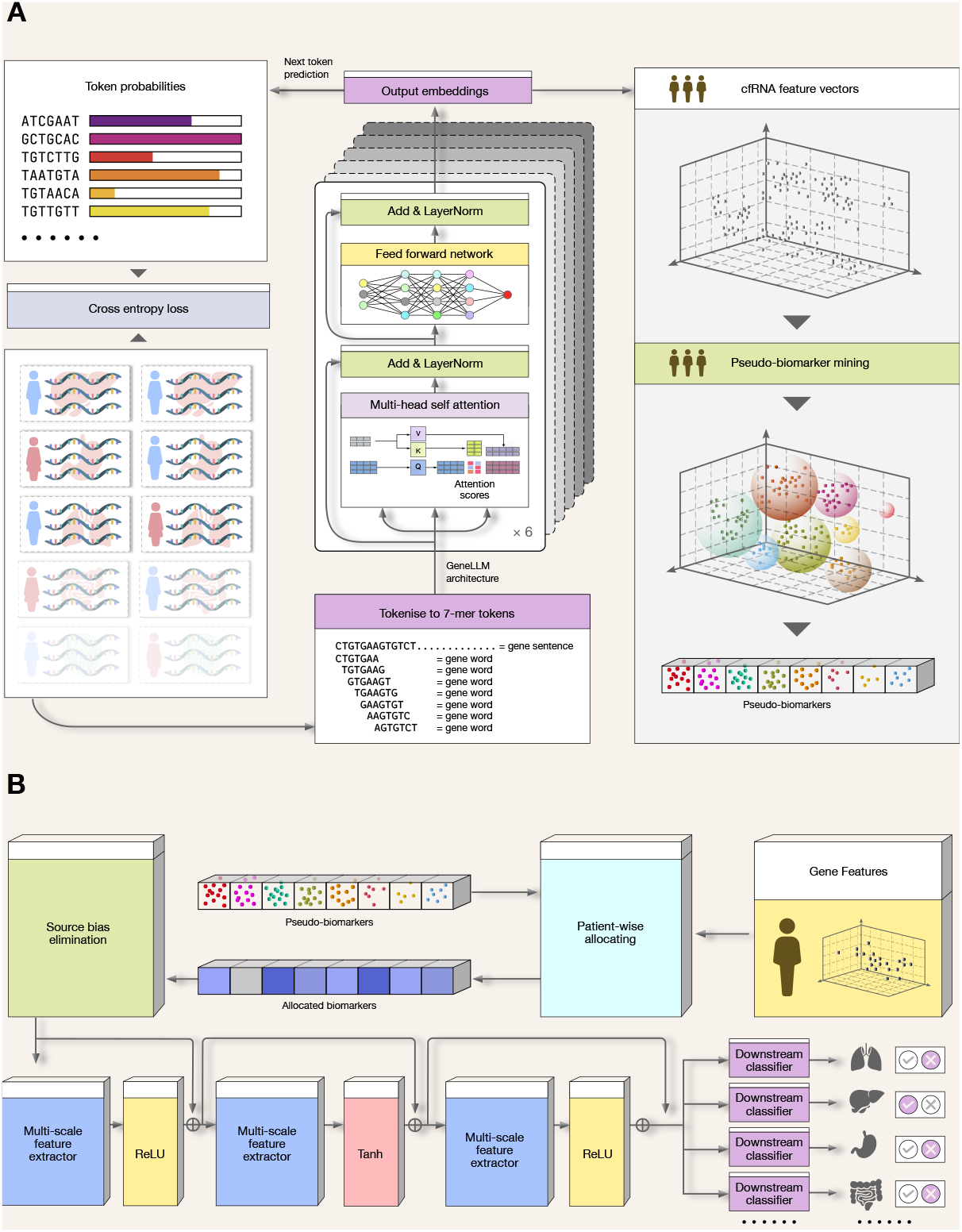
The architecture of GeneLLM. (A) The pre-training and pseudo-biomarker mining stages of GeneLLM. CfRNA reads are tokenised into 7-mer sequences and used to pretrain the large language model (LLM). Subsequently, each cfRNA read is transformed into a feature vector (upper right). These vectors are then used for mining pseudo-biomarkers. (B) Disease tuning stage of GeneLLM. Pseudo-biomarkers are used to allocate feature vectors for each patient. Multi-scale feature extractors uncover deeper information within these features. Multiple binary classifiers, one per disease, generate predictions. A source bias elimination step precedes deep feature extraction to mitigate potential bias introduced by the patient’s source hospital.

### 3.2 GeneLLM performance evaluation

A comprehensive dataset of 496 cfRNA cases, comprising both cancer patients and non-cancer individuals, was assembled for this study. To rigorously evaluate GeneLLM’s performance, the dataset was split into training, validation, and test sets at a ratio of 5 : 1 : 4, respectively. GeneLLM was pretrained exclusively on the training set, and the reported results are based on its performance on the held-out test set, ensuring an unbiased assessment of its generalisation ability.

Figures 3A and B present the Receiver Operating Characteristic (ROC) curve and confusion matrix, respectively. The ROC curve demonstrates exceptional performance, consistently exceeding 0.9 for each cancer type and pan-cancer (Figure 3A). This high Area Under the Curve (AUC) indicates GeneLLM’s strong ability to discriminate between cancer patients and non-cancer individuals. The confusion matrix (Figure 3B) further supports this, showing an average accuracy of 83.0% for classifying individual cancer types.

**Figure 3.**
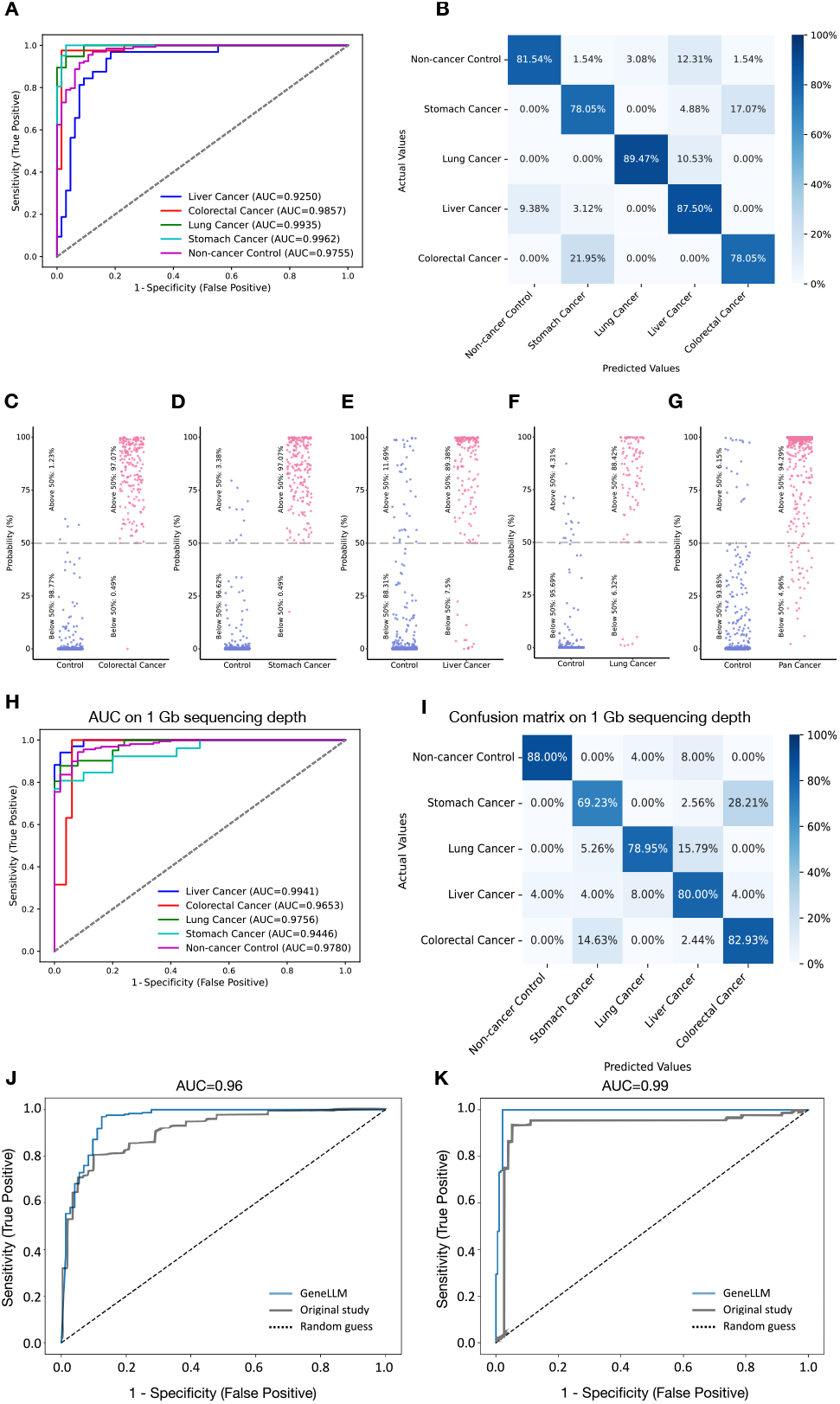
GeneLLM performance. (A) The ROC curves for four cancer types and pan-cancer, showing AUC values. (B) Confusion matrix for four cancer types and non-cancer controls. (C–G) Probability distributions calculated by GeneLLM for each sample of (C) colorectal cancer, (D) stomach cancer, (E) liver cancer, (F) lung cancer, and (G) pan-cancers with their corresponding non-cancer controls. Each dot represents a sample. (H) ROC curves for four cancers and pan-cancers using 1 GB of sequencing depth. (I) Confusion matrix for 1 GB of sequencing depth. (J) Comparison of AUC values for GeneLLM and the study by Chen *et al*. (2022) [8] using the same pan-cancer datasets. (K) Comparison of AUC values for GeneLLM and the study by Yu *et al*. (2020) [32] using the same PDAC datasets.

The predicted probabilities for each case being identified as a specific cancer type are shown in Figures 3C–G, providing a detailed view of the model’s performance at the individual case level. This further reinforces the reliability and precision of GeneLLM in identifying specific diseases from cfRNA profiles.

These results highlight the potential of GeneLLM for early cancer detection. The observed accuracy and robustness suggest its potential for clinical integration, aiding in the early detection of various cancer types and potentially contributing to improved patient outcomes.

### 3.3 GeneLLM performance with shallow sequencing depth

Standard cfRNA-seq data from plasma samples typically yields 6 GB of data, equivalent to approximately 40 million paired-end reads. We investigated GeneLLM’s robustness and versatility when using cfRNA-seq data with significantly reduced sequencing depth, mimicking real-world shallow sequencing scenarios.

To evaluate GeneLLM’s performance under these conditions, the same library was re-sequenced to generate a 1 GB dataset, representing a substantial reduction in sequencing depth. GeneLLM was then pretrained from scratch using this 1 GB dataset and subsequently fine-tuned and tested using the same 1 GB data, following the same workflow as the previous analysis. This allowed for a direct comparison of performance across different sequencing depths.

Despite the substantial reduction in sequencing depth, GeneLLM maintained its predictive accuracy. The results in Figures 3H & I demonstrate comparable performance to that observed with the higher sequencing depth data.

These findings highlight the potential applicability of GeneLLM in real-world settings where shallow sequencing is employed due to cost or resource limitations. The model’s robustness with reduced sequencing depth suggests its ability to deliver accurate predictions even under challenging conditions. Furthermore, GeneLLM’s sustained performance with limited sequencing information suggests its efficient utilisation of the most informative features within the cfRNA-seq data, which is particularly valuable when high-depth sequencing is impractical.

### 3.4 Comparison of GeneLLM with existing methods on public datasets

To benchmark GeneLLM’s performance, we compared it against published results on publicly available cfRNA datasets. These included datasets from Yu *et al*. 2020 (NCBI BioProject PRJNA552230, comprising 284 plasma cfRNA samples in extracellular vesicles (EVs) with pancreatic ductal adenocarcinoma (PDAC), 100 patients with chronic pancreatitis, and 117 healthy controls) [32], and Chen *et al*. 2022 (NCBI BioProject PRJNA729258, containing plasma cfRNA samples from 54 colorectal cancer, 37 stomach cancer, 27 liver cancer, 35 lung cancer, 31 esophageal cancer, and 46 healthy controls) [8]. Raw `fastq` data from these studies were preprocessed using our established pipeline, and GeneLLM was retrained on the processed reads. The results are presented in Figures 3J & K.

GeneLLM demonstrated improved performance in cancer detection. For pan-cancer detection, GeneLLM achieved an AUC of 0.96, outperformed to 0.91 reported by Chen *et al*. [8] (*p* < 0.05, Wilcoxon signed-rank test [15]). For pancreatic cancer detection, GeneLLM achieved an AUC of 0.99, compared to 0.95 reported by Yu *et al*. [32] (*p* < 0.05, Wilcoxon signed-rank test). These improvements underscore the efficacy of GeneLLM in enhancing cancer detection capabilities.

## 4 Validation and benchmarking on GeneLLM and pseudo-biomarkers

### 4.1 GeneLLM discovered ‘dark matter’ in the genome that aids in cancer identification

Having established the architecture and evaluated the performance of GeneLLM, we now investigate the sequences identified as key differentiators between cancerous and non-cancerous samples. This analysis aims to characterise these sequences and elucidate the mechanisms underlying GeneLLM’s discriminatory power. We focus on validating the model’s ability to leverage information from unannotated genomic regions and benchmarking its performance against established classification methods.

To determine the genomic origin of the most informative sequences identified by GeneLLM, we extracted the top three pseudo-biomarkers–short, indicative cfRNA sequences– for each cancer type. A Nucleotide Basic Local Alignment Search Tool (BLASTn) search against the NCBI nucleotide collection (nt) database (updated 23rd June 2024) [6] revealed that most pseudo-biomarkers did not align with any annotated genes (Figure 4), despite high coverage and identity (> 95% and > 99%, respectively) when mapped to the human genome. This suggests that GeneLLM leverages information from unannotated genomic regions–often termed ‘dark matter’–for cancer identification. This capability distinguishes GeneLLM from traditional gene expression-based methods, which are limited by their reliance on existing annotations. Further investigation into these unannotated regions may yield insights into cancer development and progression [14, 18, 22].

**Figure 4.**
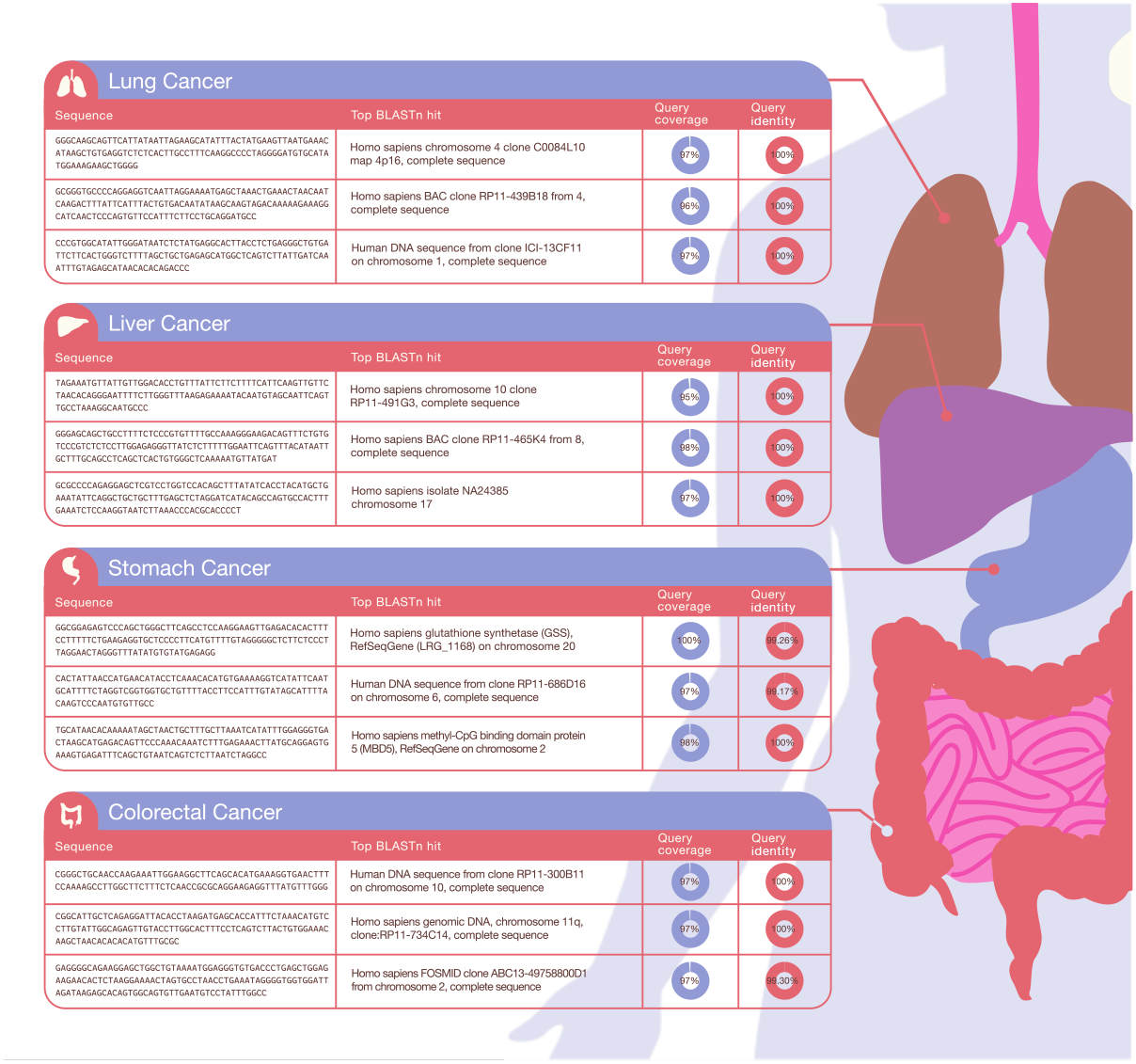
Top three pseudo-biomarkers for each cancer types and their corresponding BLASTn top hits.

To assess pseudo-biomarker specificity, we examined the scores (The method to calculate the scores are illustrated in Section 6.6.3) assigned by GeneLLM to the top three pseudo-biomarkers for each cancer type across all samples (Figure 5). We observed a clear association between specific pseudo-biomarkers and their corresponding cancer types. For example, pseudo-biomarkers identified for stomach cancer exhibited high scores in stomach cancer samples but significantly lower scores in other samples. This pattern, consistently observed across all cancer types, indicates that GeneLLM identifies pseudo-biomarkers with high cancer-type specificity, underscoring their potential as diagnostic tools.

**Figure 5.**
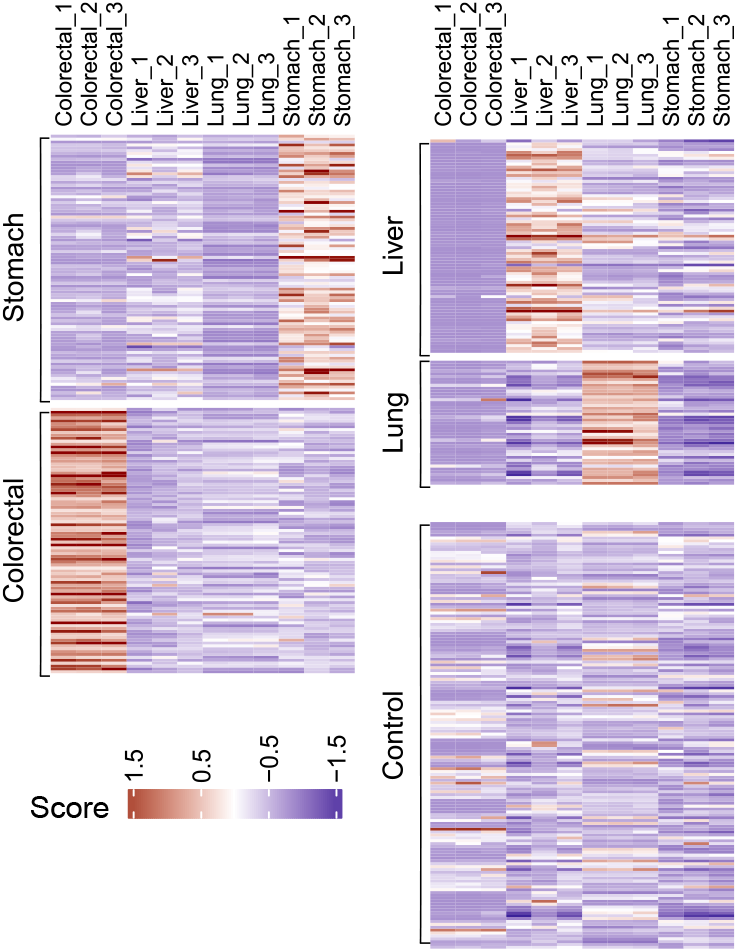
Scores assigned to the top three pseudobiomarkers for each cancer type.

Finally, we benchmarked GeneLLM’s performance against established machine learning methods, including support vector machine, random forest, XGBoost, logistic regression, elastic net, and a plain neural network (Figure 6). Using normalised expression matrices derived from the same cfRNA sequencing data, GeneLLM consistently outperformed all other methods across various performance metrics. This superior performance, combined with its ability to leverage information from unannotated genomic regions, positions GeneLLM as a powerful tool for cancer detection and biomarker discovery.

**Figure 6.**
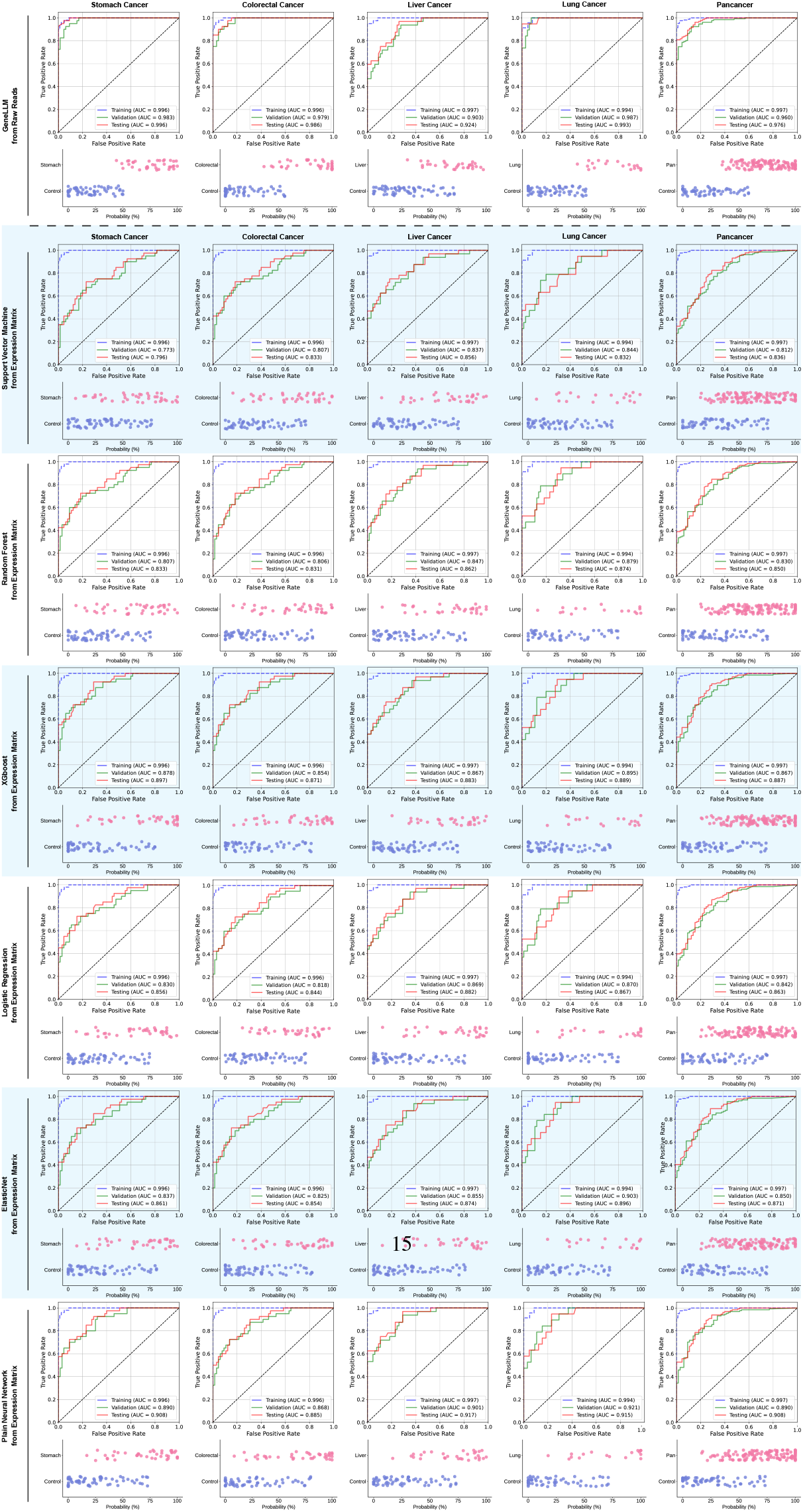
Benchmarking GeneLLM performance from raw reads against other classification methods using normalised expression matrices.

## 5 Discussion

Applying GeneLLM to cancer detection represents a significant advance in leveraging information encoded within cfRNA. The model’s high performance, demonstrated by AUC values exceeding 0.9 across multiple cancer types and consistent accuracy in classifying individual cancers, suggests a robust and reliable approach for early detection. This stems from GeneLLM’s ability to capture subtle, complex patterns within cfRNA data—patterns often overlooked by traditional methods relying on differential expression analysis of known genes. Analogous to language model training [25], the pretraining strategy allows GeneLLM to learn intrinsic relationships between cfRNA sequences, enabling it to discern minute, disease-indicative variations. This inherent ability to learn contextual information directly from raw sequencing data distinguishes GeneLLM from annotation-dependent methods.

GeneLLM’s maintained high performance even with significantly reduced sequencing depth (one-sixth of standard depth) is particularly noteworthy. This robustness not only reduces costs and computational burden but also suggests efficient identification and utilisation of the most informative cfRNA features. This efficiency may derive from the model’s focus on short, indicative sequences (‘pseudo-biomarkers’) rather than entire gene expression levels. This targeted approach is advantageous when cfRNA is fragmented or at low concentrations, common challenges in liquid biopsy diagnostics. Further investigation into pseudo-biomarker characteristics (length distribution, sequence composition, and genomic origin) could provide valuable insights into cancer biology and inform the development of more targeted diagnostic tools. These pseudo-biomarkers may represent fragments of larger transcripts from unannotated regions or arise from novel, cancer-specific transcriptional events.

The discovery that many pseudo-biomarkers originate from unannotated genomic regions (‘dark matter’) has profound implications. This suggests that GeneLLM accesses a previously unexplored source of diagnostically relevant biological information. The high conservation of these unannotated regions across samples (high coverage and identity in BLASTn searches), despite their lack of association with known genes, raises intriguing questions about their function and role in cancer development. Unravelling the mysteries of these unannotated regions could open new avenues for understanding cancer biology and developing novel therapeutic strategies. GeneLLM’s ability to identify these sequences highlights the potential of AI-driven approaches to uncover hidden biological signals and accelerate scientific discovery.

The specificity of the identified pseudo-biomarkers, demonstrated by their distinct scoring patterns across different cancer types, further strengthens their diagnostic potential. This specificity suggests that different cancers may exhibit unique cfRNA signatures derived from these unannotated regions, potentially reflecting distinct molecular mechanisms driving their development and progression. This observation raises the possibility of developing highly specific diagnostic tests tailored to individual cancer types, enabling earlier and more accurate diagnosis [19, 23]. Moreover, GeneLLM’s superior performance compared to traditional machine learning methods trained on annotated gene expression data underscores its ability to extract and utilize more informative features from cfRNA. This suggests that the information within these unannotated regions, combined with GeneLLM’s pattern recognition capabilities, provides a more comprehensive and accurate understanding of the underlying disease state.

Comparison of GeneLLM with existing methods using public datasets further validates its superior performance. The improved AUC values achieved by GeneLLM for both pancancer and pancreatic cancer detection, compared to previously published results [8, 32], demonstrate its enhanced ability to detect cancer using existing cfRNA data. This improved performance, coupled with reduced sequencing depth requirements, positions GeneLLM as a cost-effective and highly accurate approach for early cancer detection. Consistent performance across multiple datasets suggests that the model’s ability to identify informative cfRNA patterns is not limited to specific cohorts or experimental conditions, further supporting its generalisability and potential clinical application. However, limitations of cross-species and cross-disease stage comparisons must be acknowledged. Further validation using larger, more diverse cohorts, including different cancer stages and diverse ethnic backgrounds, is crucial to fully assess clinical utility and ensure equitable performance across populations.

The implications extend beyond cancer diagnostics. GeneLLM’s ability to analyse raw sequencing data and identify informative sequences without relying on existing annotations opens new possibilities for biomarker discovery in other diseases and biological contexts. The framework is adaptable to other sequencing data types, such as intracellular bulk RNA-seq, single-cell RNA-seq, or even DNA sequencing data, to identify novel biomarkers or therapeutic targets. For example, applying GeneLLM to single-cell RNA-seq data could reveal cell-type-specific pseudo-biomarkers associated with disease progression or treatment response. Similarly, analysing DNA sequencing data could uncover previously unknown disease-associated genetic variations or structural alterations. The potential applications of this technology are vast and could revolutionise our understanding of complex biological processes and accelerate the development of personalised medicine.

While GeneLLM demonstrates remarkable potential, potential limitations and future research directions warrant attention. Biological characterisation of the identified pseudo-biomarkers is crucial. Determining their origin and function is essential for understanding the mechanisms driving their association with cancer. This could involve experimental validation using techniques such as quantitative PCR, *in situ* hybridisation, or functional assays. Investigating the roles of these sequences in cancer development and progression could lead to the identification of novel therapeutic targets. The potential impact of technical artefacts or biases in the sequencing data on GeneLLM’s performance requires careful evaluation of the model’s robustness to variations in sequencing platforms, library preparation methods, and data processing pipelines to ensure reliable and reproducible results. Finally, integrating GeneLLM with other data modalities (clinical, imaging, or proteomic data) could further enhance diagnostic accuracy and provide a more holistic view of the disease state. Addressing these limitations and pursuing these future research directions can unlock GeneLLM’s full potential and pave the way for a new era of precision medicine.

## 6 Methods

### 6.1 Ethics approval

The ethics of this study were approved by the Ethics Committee and the Institutional Review Board of Peking University First Hospital (2022-132), Beijing Hospital (2022BJYYEC-420-02), and Sir Run-Run Shaw Hospital (2023-0300). Written informed consent was obtained from all patients prior to participation.

### 6.2 Cohort design

The cohort for this study comprised 496 plasma samples, including 102 colorectal cancer and 102 stomach cancer samples from Peking University First Hospital, 81 liver cancer samples from Sir Run-Run Shaw Hospital, 47 lung cancer samples from Beijing Hospital, and 164 non-cancer controls. The non-cancer controls were obtained from all three centres to calibrate potential batch effects.

### 6.3 Sample collection

Peripheral whole blood samples were taken from subjects before treatment using EDTA-coated tubes. The tubes were inverted between 8 and 10 times to ensure the mixing of the blood with the anticoagulant. Plasma separation was performed following a standard clinical protocol for blood centrifugation within two hours of blood collection. After separation, the plasma samples were divided into aliquots and preserved at –80°C until the extraction of cfRNA.

### 6.4 CfRNA extraction, library preparation, and sequencing

Total cfRNA was isolated from 200 *μ*L of plasma utilising the miRNeasy Serum/Plasma Advanced Kit (Qiagen), which employs Qiagen’s unique resin as the separation matrix. This kit is capable of extracting all RNA molecules from approximately 18 nt upwards. DNase I (NEB) was used to eliminate remaining DNA.

To assess potential contamination in the RNA extraction and library preparation processes, two types of negative controls were utilised. These included libraries generated from RNA/DNA-free water (Invitrogen) and solutions eluted from the miRNeasy Serum/Plasma Advanced Kit with RNA/DNA-free water as input. A non-reverse transcriptase control was implemented to prevent DNA contamination. The positive control comprised human brain RNA, provided SMARTer Stranded Total RNA-Seq Kit, and was processed using the same protocol as that for the cfRNA library preparation.

The comprehensive cfRNA library (approximately > 50 nt) was constructed using the SMARTer Stranded Total RNA-Seq Kit v2-Pico Input Mammalian (Takara), adhering to the manufacturer’s recommended protocol. This kit employs a CRISPR/DASH technique to eliminate ribosomal cDNA. Each library was sequenced over 40 million reads using Illumina NovaSeq6000 platform using a PE150 strategy.

### 6.5 Data quality control and preprocessing

To ensure high-quality data, we employed strict quality control processes. First, all raw data were trimmed using fastp v0.23.3 [7] with default parameters to automatically trim low-quality reads and adapters. The trimmed reads were then mapped to the human genome (GENCODE human release 43 [13]) using STAR v2.7.10b [11] with the encyclopedia of DNA elements (ENCODE) standard options [12, 16]. The mapped reads were used as input for GeneLLM. Mapping statistics were then retrieved using the flagstat function in SAMtools v1.17 [10].

### 6.6 AI model

#### 6.6.1 Inputs and data sources

The input for GeneLLM is a sequence of tokens derived from the cfRNA sequencing reads of both cancer patients and individuals without cancer. Each patient contributes approximately 40 million cfRNA reads. During the pretraining phase, GeneLLM focuses on understanding the internal relationships within the cfRNA reads and storing pertinent information without using cancer labels. These cfRNA reads are then converted into 7-mers, each considered a token.

For batch processing, the lengths of different ID sequences are standardised by padding, inserting [PAD] tokens to ensure all token sequences in a batch are of equal length. Consistent with Natural Language Processing (NLP) tasks, specific tokens [SOS] and [EOS] are employed to mark the start and end of each sequence, respectively.

#### 6.6.2 Pretraining

During the pretraining phase described in Figure 2A, each cfRNA sequence is treated as an independent sentence, with 7-mer sequences acting as tokens. The core model architecture is a deep Transformer decoder network. To distinguish between input and target sequences during training, a special [SOS] (start of sequence) token is prepended to the input tokens, and an [EOS] (end of sequence) token is appended to the target tokens. The model employs a standard language modelling objective, learning to predict the next token in the target sequence based on the input sequence and preceding target tokens.

The model is composed of six Transformer blocks stacked sequentially. Each block comprises a self-attention sublayer and a feed-forward sublayer, adhering to the original Transformer architecture proposed by Vaswani *et al*. (2017) [31]. A triangular masking pattern is applied to the self-attention matrices within each block. This masking enforces the autoregressive property of the decoder, ensuring predictions for a given target token to depend only on previous tokens in the sequence and the input tokens, never on future tokens.

Pretraining on unlabelled cfRNA sequencing data enables the model to learn meaningful representations that can be fine-tuned for downstream tasks.

#### 6.6.3 Pseudo-biomarker mining

Once the model is pretrained, we fix the parameters and use it to mine hidden information within the massive data of human cfRNA reads. Since each patient typically has approximately 40 million cfRNA reads, we first use GeneLLM to encode each read into an embedding. Denote the *i*-th patient’s *j*-th cfRNA read as *s*_*ij*_, we transformed it into a gene feature vector *g*_*ij*_ as shown in Equation (1), with an illustrative process in Figure 2A.

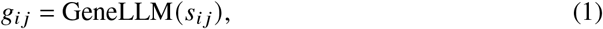

where *g*_*ij*_ ∈ 𝒢 is the last output vector.

For accurate and interpretable cancer detection, we need to identify ‘biomarkers’, referred to as ‘pseudo-biomarkers’ in this study. These pseudo-biomarkers are vectors that best represent all cfRNA vectors *g*_*ij*_. Denoting the pseudo-biomarker vectors as *ν*_1_, …, *ν*_*n*_, each cfRNA representation *g*_*ij*_ should match a pseudo-biomarker that best represents it. Therefore, we aim to minimise Equation (2).

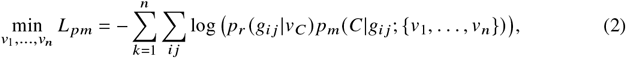

where *C* represents an index of the pseudo-biomarkers, and *p*_*m*_ represents the decision probability for each cfRNA vector, defined as:

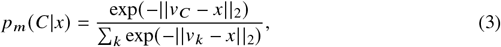

where *p*_*r*_ represents the reconstruction probability.

Inspired by Sha *et al*. (2021) [28], each pseudo-biomarker is defined as a Gaussian distribution, with *ν*_*C*_ as the mean and *σ*_*C*_ as the standard deviation. The reconstruction probability is calculated by:

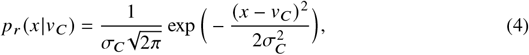

where *C* is the index for pseudo-biomarkers, and both *ν*_*C*_ and *σ*_*C*_ are trainable parameters. This pseudo-biomarker mining phase can be seen as an Large Language Model (LLM) alignment process, aligning the information in GeneLLM with the pseudo-biomarkers by minimising *L* _*pm*_ in Equation (2). After this process, the pseudo-biomarker vectors *ν*_1_, …, *ν*_*n*_ are used to extract important features from the extensive cfRNA reads for each patient.

##### Pseudo-biomarker scoring

To investigate the relationship between pseudo-biomarkers and diseases, we first identified the three most indicative cfRNA sequences (pseudo-biomarkers) for each cancer type. For cancer type *d*_*k*_, let these pseudo-biomarkers be *ν* (1), *ν* (2), and *ν*(3). Their corresponding representations, 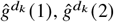, and 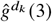, were calculated as:

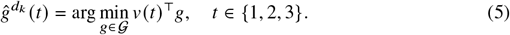

For patient *i*, the score associated with each indicative pseudo-biomarker is:

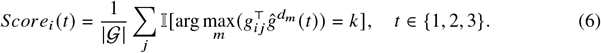

which quantifies the proportion of patient *i*’s cfRNA reads that align with the corresponding indicative pseudo-biomarker.

#### 6.6.4 Disease tuning

Since each patient has approximately 40 million cfRNA reads, summarising these features is essential. This is achieved using pseudo-biomarkers, as illustrated in Figure 2B. For the *i*-th patient, we summarise the feature as follows:

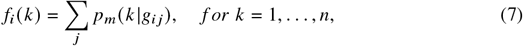

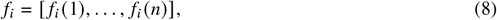

where *f*_*i*_ (*k*) represents the feature for pseudo-biomarker vector *ν*_*k*_.

##### Source bias elimination

To address biases from different sources (centres), we train a mapping for each patient source to align the non-cancer cases from each source to a common distribution. This process is depicted in Figure 2B as the source bias elimination module.

Formally, assuming the *i*-th patient has a feature vector *f*_*i*_ and a source *s*_*i*_, we collect a set of all healthy cases from the training set, denoted as ℋ. We then calculate the midpoint for each source:

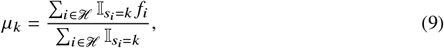

where *μ*_*k*_ represents the midpoint value for the *k*-th source, with 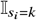 as an indicator function, which is 1 when *s*_*i*_ equals to *k*, and 0 otherwise. Next, we align all the midpoints to one standard midpoint, *μ*_*_^1^. Each patient’s feature vector is then corrected as follows:

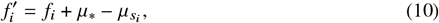

where 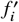 is the corrected feature vector without source bias.

##### Multi-scale feature extractor

The feature vector, derived from the vast amount of cfRNA reads, exhibits a rich set of multi-scale features that capture intricate patterns and variations at different levels of granularity. These multi-scale features are crucial for understanding the complex nature of the cfRNA sequences and their potential role in disease detection.

Inspired by the Dense Convolutional Network architecture proposed by [17], we propose to extract multi-scale features by employing stacked skip-connection layers. As illustrated in Figure 2B, we utilize multiple Multi-scale Feature Extractor modules with activation functions such as the ReLu(·) [1] and the Tanh(·) to extract multi-scale features. Within each module, we iteratively apply a feedforward layer to the input feature vector and concatenate the output to the input, effectively preserving information from previous layers and enabling the network to learn multi-scale representations.

The proposed architecture leverages the power of skip connections, which allow the network to propagate information from earlier layers to later layers, mitigating the vanishing gradient problem and facilitating the learning of multi-scale features. By iteratively applying feedforward layers and concatenating the outputs, the network can capture a wide range of patterns and variations present in the cfRNA sequences at different scales. This approach enables the model to effectively learn and represent the complex characteristics of the cfRNA reads, which is essential for accurate disease detection.

By incorporating multiple Multi-scale Feature Extractor modules, the proposed architecture can extract a comprehensive set of multi-scale features that encapsulate the rich information present in the cfRNA sequences. These features serve as a robust foundation for subsequent analysis and disease detection tasks, enabling the development of more accurate and reliable diagnostic tools based on cfRNA sequencing data.

##### Multiple comparable downstream classifier head

In this part, we aim to use each patient’s feature vector to detect any possible diseases they may have. Given the multi-scale feature vector for the *i*-th patient, denoted as 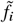, we assume that we are going to detect *N*_*d*_ types of diseases. We construct a feed-forward classifier for each disease as follows:

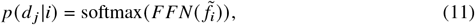

where ‘FFN’ represents the feed-forward layer, and *d* _*j*_ represents the *j*-th disease. Each classifier head is trained using samples from patients with that specific disease and non-cancer controls.

However, there is still one issue: the probabilities calculated by different classifiers usually cannot be directly compared. For example, a patient may have a probability of 0.6 according to the stomach cancer classifier and 0.5 according to the lung cancer classifier. However, if the threshold for the stomach cancer classifier is 0.7 and the threshold for the lung cancer classifier is 0.4, this patient should be classified as having lung cancer. If we simply compare the absolute values of the probabilities, we will obtain an incorrect result. Therefore, it is necessary to calibrate the disease screening probabilities across all classifiers. Here, we use a simple yet effective method called ‘pivot calibration’. Given the threshold 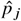 for the *j*-th disease and a general threshold *p*_*M*_, we force every 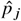 to be equal to *p*_*M*_, and every probability is tuned by ratio. The detailed calibration process is described in Eqn. (12).

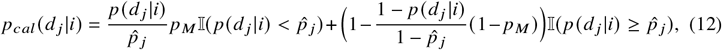

where 𝕀 (·) represents the indicator function, which equals 1 if the condition inside the parentheses is true and 0 otherwise.

Finally, we compare the calibrated probability *p*_*cal*_ (*d* _*j*_|*i*) of every disease and select the disease with the highest probability as our prediction. If every probability is below *p*_*M*_, then the patient is classified as non-cancer.

#### 6.6.5 Training regimen

To train the GeneLLM, cfRNA reads were preprocessed as described in Section 6.6.1.

We trained the GeneLLM on 64 NVIDIA A100 GPUs, utilising their high-performance computing capabilities. To optimise memory usage and training efficiency, we used a batch size of 1 per GPU. The model was pre-trained using the entire training set of cfRNA sequences, allowing it to learn the underlying patterns and characteristics of the data. The pre-training process was computationally intensive and took approximately 15 days to complete.

After pre-training, we utilised the entire cfRNA dataset to extract pseudo-biomarkers. To further refine the GeneLLM for disease prediction, we performed disease tuning using labelled patient cfRNA sequential data. Each patient in the dataset has a massive amount of cfRNA sequences associated with their condition. By leveraging this labelled data, we fine-tuned the pre-trained GeneLLM to capture disease-specific patterns and characteristics. The disease tuning process adapts the model to better discriminate between different disease states and improves its performance in downstream tasks such as disease prediction.

The combination of pre-training on a large cfRNA dataset and subsequent disease tuning using labeled patient data enables the GeneLLM to learn both general and disease-specific representations of cfRNA sequences. This approach harnesses the power of transfer learning, where the knowledge gained from pre-training on a diverse dataset is transferred and specialised for specific disease applications through fine-tuning. The resulting model is expected to capture relevant biomarkers and patterns associated with various diseases, potentially enhancing the accuracy and efficiency of disease diagnosis and monitoring using cfRNA data.

#### 6.6.6 Inference regimen

During the inference process, we sequentially input the cfRNA sequences into the GeneLLM, followed by the disease judgment head. The GeneLLM encodes each cfRNA sequence, capturing its relevant features and representations. The encoding time varies depending on the length of the cfRNA sequences, as longer sequences require more computational processing. On average, the GeneLLM takes approximately 2.16 × 10^−5^ seconds to encode a single cfRNA sequence. However, it’s important to note that each patient typically has a massive amount of cfRNA sequences associated with their sample. Therefore, the total encoding time for a single patient is around 14.4 minutes, considering the processing of all their cfRNA sequences.

After encoding the cfRNA sequences, the disease judgement head comes into play. This component of the model is responsible for making predictions or classifications regarding the presence or absence of specific diseases based on the encoded representations. To enhance the robustness and reliability of the disease judgement, we employ an ensemble approach. We use 10 ensemble models, each trained on different subsets or variations of the data, to make predictions. The ensemble approach helps to mitigate potential biases or variances in individual models and improves the overall accuracy of the disease judgement.

The inference time for the disease judgement head, using the 10 ensemble models, takes approximately 5.6 minutes on average for one patient. This inference time includes the processing of the encoded cfRNA sequences by each ensemble model and the aggregation of their predictions to reach a final judgement.

Compared to previous methods for cancer screening, the combination of the GeneLLM and the disease judgement head offers a remarkably quick and efficient approach. Traditional cancer screening methods often involve invasive procedures, extensive laboratory tests, and manual analysis, which can be time-consuming and resource-intensive. In contrast, our proposed method leverages the power of machine learning and natural language processing techniques to analyze cfRNA sequences rapidly and accurately.

It’s worth noting that while the inference times mentioned are impressive, they are based on the specific hardware and computational resources used in our experiments. The actual inference times may vary depending on the available computational infrastructure and the scale of the patient cohort being analyzed. Nevertheless, the efficiency and speed of our method highlight its potential for widespread adoption in clinical settings, enabling large-scale cancer screening programs and improving patient care.

#### 6.6.7 Metrics

To evaluate the architecture’s performance in detecting diseases, we calculated the confusion matrix and Area Under the Receiver Operating Characteristic Curve (AUC-ROC). The confusion matrix provides a tabular summary of the model’s classification results, displaying the counts of true positive (TP), true negative (TN), false positive (FP), and false negative (FN) predictions. These counts offer insights into the model’s ability to correctly identify positive and negative samples and the types of errors it makes. The AUC-ROC value represents the probability that a randomly selected positive sample will have a higher score assigned by the model than a randomly selected negative sample. A larger AUC value, ranging from 0 to 1, suggests better model performance in distinguishing between positive and negative samples. Furthermore, we plotted the ROC curve to visually assess the model’s performance. The ROC curve illustrates the trade-off between the true positive rate (TPR) and the false positive rate (FPR) at various classification threshold values. An ideal ROC curve would hug the top-left corner of the plot, indicating high TPR and low FPR across a wide range of thresholds. The closer the ROC curve is to the top-left corner, the better the model’s performance. The AUC value summarises the overall performance of the model across all possible thresholds, providing a single scalar value for comparison. By evaluating the architecture’s performance using these metrics and visualisations, we gain a comprehensive understanding of its effectiveness in detecting diseases.

## Supplementary information

**Table S1. Patient information**.

## Declarations

### Consent to participate

Participants in this study were fully informed about the aims, procedures, and potential risks involved. All participants provided written informed consent before participation, and the study followed ethical guidelines consistent with the 1964 Helsinki declaration and its later amendments. Additionally, the study protocol was reviewed and approved by the Ethics Committee and the Institutional Review Board of Peking University First Hospital, Beijing Hospital, and Sir Run-Run Shaw Hospital, under approval number 2022-132, 2022BJYYEC-420-02, and 2023-0300, respectively.

### Consent for publication

The authors affirm that all participants understand that their anonymised data will be published in a scientific journal. Consent for publication was also obtained from all participants at the time of consent for participation.

### Availability of data and materials

All the raw sequencing data (including shallow sequencing) have been deposited in the National Center for Biotechnology Information (NCBI) Sequence Read Archive (SRA) database under the BioProject accession PRJNA1128376.

### Code availability

The source code of GeneLLM, the model weights, and the inference code will be released after the paper is published.

## Authors’ contributions

The methodology and algorithm of GeneLLM, validated in this study, had been fully developed by L.S. with support from and exclusive authorisation granted to OxTium Technology Co., Ltd. S.D., Y.J., T.Z. and L.S. conceived current project idea. S.C. and P.W. designed clinical validation plan in this article. Y.J. designed and carried out wetlab experiments. S.D. designed and performed all bioinformatics. L.S. conceived, wrote and optimised artificial intelligent algorithms. T.Z., S.C., J.X., C.W. and P.W. coordinated human samples. Q.L. improved artificial intelligence. C.X., Y.X., Y.L., M.H. and Y.G. carried out various experiments. S.D., L.S., Y.J, T.Z., and S.C. wrote and edited manuscript. All authors discussed the results and commented on the manuscript.

## Acknowledgments

This work was financially supported by the National Key R&D Program of China (No. 2022ZD0117700), Shenzhen Science and Technology Program (No. 20240724152335001), the National Natural Science Foundation of China (No. KZ37117501, No. ZG216S23E8, No. 62306024, and No. 82271767), the Beijing Hospital New-Star Plan of Science and Technology (BJ-2020-084), the Startup Foundation for Doctors of Beijing Hospital (No. BJ-2019-132), and Capital’s Funds for Health Improvement and Research (No. 2022-2-4075 and No. 2024-4-40711). We also recognise the investors in OxTium Technology, whose commitment to advancing the boundaries of technology has been invaluable. We acknowledge the assistance of Jian Guo, Yelin Lai, Junhao Chen, and Yunlong Hao in this study.

## Footnotes

1 In practice, the *μ*_*_ can be set as the average value of all midpoints.

